# Untargeted metabolomic profiling reveals mTORC1-dependent regulation of amino acid utilization in lymphatic endothelial cells

**DOI:** 10.64898/2026.02.17.706183

**Authors:** Jie Zhu, Felix Darko, Fei Han, Summer Simeroth, Lingjun Li, Haiwei Gu, Pengchun Yu

**Author notes:** These authors contributed equally: Jie Zhu and Felix Darko. Correspondence: Pengchun Yu, Ph.D. Cardiovascular Biology Research Program Oklahoma Medical Research Foundation, 825 NE 13^th^ Street, Oklahoma City, OK 73104.

## Abstract

The lymphatic vascular system plays essential roles in tissue fluid drainage, dietary fat absorption and transport, and immune cell trafficking. To support these physiological functions, the lymphatic vasculature forms an extensive and highly organized network throughout the body. We have recently discovered that the mechanistic target of rapamycin complex 1 (mTORC1), with RAPTOR as an indispensable component, directs glycolysis and glutaminolysis in lymphatic endothelial cells (LECs) to promote lymphatic vessel formation. However, the role of mTORC1 in regulating LEC metabolism remains incompletely understood. Here, by conducting untargeted metabolomic profiling of control and RAPTOR-deficient LECs, we uncover a global impact of mTORC1 inhibition on amino acid utilization. Specifically, RAPTOR deficiency impairs the conversion of glutamine to glutamic acid, resulting in decreased levels of glutamic acid and aspartic acid, as well as reduced abundance of N-acetyl-glutamic acid and N-acetyl-aspartic acid—two metabolites unexpectedly detected in LECs. Integrated metabolomic and transcriptomic analyses further reveal that impaired glutaminolysis in RAPTOR-depleted LECs is accompanied by an increase in intracellular asparagine, arginine, and metabolites associated with arginine catabolism, potentially driven by upregulation of their respective transporters. In addition, RAPTOR depletion results in abnormal accumulation of branched-chain amino acids (BCAAs) and other essential amino acids primarily involved in protein synthesis. Mechanistically, our data suggest that defective BCAA catabolism and impaired translational control contribute to these metabolic alterations. Collectively, these findings reveal an important role of mTORC1 signaling in coordinating amino acid utilization and suggest that this regulation is critical for lymphatic vessel formation.

## INTRODUCTION

The lymphatic vascular system, composed of lymphatic capillaries and collecting lymphatic vessels, plays a key role in maintaining tissue fluid homeostasis by absorbing excess interstitial fluid and returning it to the venous circulation.^1^ In addition, the lymphatic vasculature transports proteins, dietary fats, cholesterol, and immune cells.^2,3^ Consequently, lymphatic vessels are implicated in several pathological conditions, including lymphedema, atherosclerosis, and inflammation.^2,3^

In order to support a wide range of physiological functions, the lymphatic vasculature forms an extensive network throughout the body.^4^ This process, known as lymphangiogenesis, begins during embryonic development and is driven by multiple growth factors, including vascular endothelial growth factor C and fibroblast growth factors.^4,5^ These ligands exert pro-lymphangiogenic effects by activating their cognate receptors and interacting with neuropilin 2, β1 integrins, ephrin B2, and other modulators of lymphatic development.^4,5^ Recent studies have also identified cellular metabolism as a key process underlying lymphangiogenesis.^6,7^ For example, we discovered that hexokinase 2 (HK2)-driven glycolysis regulates the formation of the primitive lymphatic plexus in mouse embryos.^8^ In addition to glycolysis, fatty acid β-oxidation, ketone body oxidation, and mitochondrial respiration have also been shown to contribute to the regulation of lymphangiogenesis.^9–13^ Collectively, these metabolic pathways provide energy and biosynthetic precursors to support metabolically demanding cellular processes during lymphatic formation, such as proliferation and migration of lymphatic endothelial cells (LECs).^6,7^

The mechanistic target of rapamycin (mTOR) is a serine/threonine protein kinase.^14,15^ Through interactions with distinct binding partners, mTOR forms two protein complexes, mTOR complex 1 (mTORC1) and mTOR complex 2 (mTORC2), which phosphorylate different downstream substrates to regulate a wide range of developmental and cellular processes.^14,15^ Previous studies have shown that mTORC1 and mTORC2 are both essential for lymphatic valve development.^16,17^ Moreover, we demonstrated that mTORC1 signaling promotes lymphatic capillary growth and collecting lymphatic vessel differentiation.^18^ Mechanistically, mTORC1 regulates glycolytic metabolism and glutaminolysis in LECs.^18^ Genetic inhibition of both metabolic pathways in LECs recapitulates the lymphatic developmental defects caused by mTORC1 inhibition.^18^ Despite our recent findings, the role of mTORC1 in coordinating metabolic programs in LECs remains incompletely understood.

To address this gap in knowledge, we performed unbiased metabolomic profiling to identify metabolites significantly altered by lymphatic endothelial mTORC1 inhibition. Through this approach, we uncovered a critical role for mTORC1 in regulating amino acid utilization in LECs.

## RESULTS

### Untargeted metabolomic analysis of control and RAPTOR-deficient LECs

To gain a comprehensive understanding of the role of mTORC1 signaling in regulating LEC metabolism, we depleted RAPTOR, which is an indispensable component of mTORC1, in human dermal LECs (HDLECs) using siRNA and conducted untargeted metabolomics to determine the impact of RAPTOR deficiency on cellular metabolism. Western blot analysis demonstrated efficient RAPTOR protein reduction following siRNA-mediated knockdown (**Fig. 1A and 1B**). Using liquid chromatography-mass spectrometry (LC-MS), we identified and quantified 513 metabolites that exhibited a coefficient of variation (CV) of less than 20% in quality control samples and were therefore included in subsequent analyses. Principal component analysis (PCA), an unsupervised dimensionality reduction method used to identify clustering patterns in complex metabolomic datasets, revealed clear separation between control HDLECs (n = 3 samples) and RAPTOR-deficient HDLECs (n = 4 samples) (**Fig. 1C**), indicating substantial metabolic differences between the two experimental groups.

**Figure 1.**
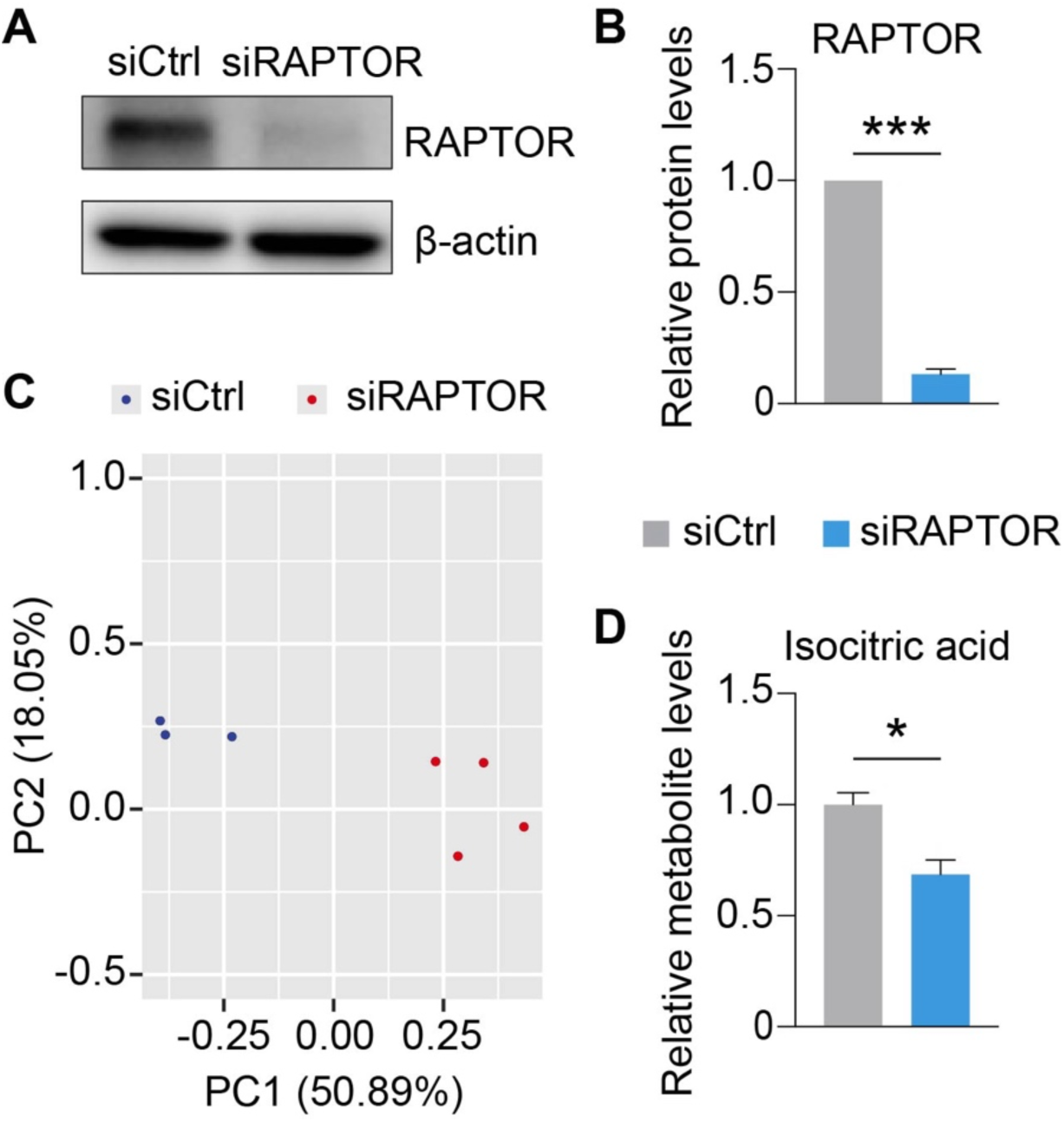
Untargeted metabolomic analysis of control and RAPTOR-deficient LECs. (A, B) Western blot analysis and densitometric quantification of RAPTOR proteins (n = 4 samples). (C) PCA plot revealing distinct metabolic profiles between control (n = 3 samples) and RAPTOR-deficient HDLECs (n = 4 samples). (D) Isocitric acid levels in control (n = 3 samples) and RAPTOR-deficient (n = 4 samples) HDLECs. Data represent mean ± SEM, *p < 0.05, ***p < 0.001, Welch’s t-test.

mTOR signaling acts as a key regulator of mitochondrial metabolism.^14,19^ Consistent with this, our recent study demonstrated that RAPTOR depletion leads to downregulation of α-ketoglutarate, succinate, fumarate, and malate in HDLECs.^18^ Therefore, to validate this untargeted metabolomics dataset, we examined whether any tricarboxylic acid (TCA) cycle metabolites were decreased in RAPTOR-deficient HDLECs, and we found that isocitric acid levels were reduced by approximately 30% (**Fig. 1D**).

### RAPTOR deficiency impairs the generation of glutamine metabolism-derived metabolites in LECs

Through statistical analysis, we identified 192 metabolites that differed significantly between control and RAPTOR knockdown HDLECs (adjusted p < 0.05). Notably, approximately 20% of these metabolites were related to amino acid metabolism, highlighting a substantial impact of RAPTOR deficiency on amino acid homeostasis. Given this strong representation of amino acid alterations, we focused our subsequent analyses on specific amino acids to determine how RAPTOR depletion affects their metabolism.

Glutamine is a crucial amino acid that fuels the TCA cycle through glutaminolysis in both blood endothelial cells and LECs, thereby impacting angiogenesis and lymphangiogenesis, respectively.^18,20–22^ We have previously shown that mTORC1 signaling in LECs promotes the expression of glutaminase (GLS), the first enzyme in glutaminolysis, which converts glutamine to glutamic acid.^18^ In line with this finding, our untargeted metabolomic profiling showed that RAPTOR depletion leads to substantial accumulation of glutamine, accompanied by significant reductions in glutamic acid and aspartic acid (**Fig. 2A–2D**), indicating a metabolic blockade at the glutamine-to-glutamic acid conversion step.

**Figure 2.**
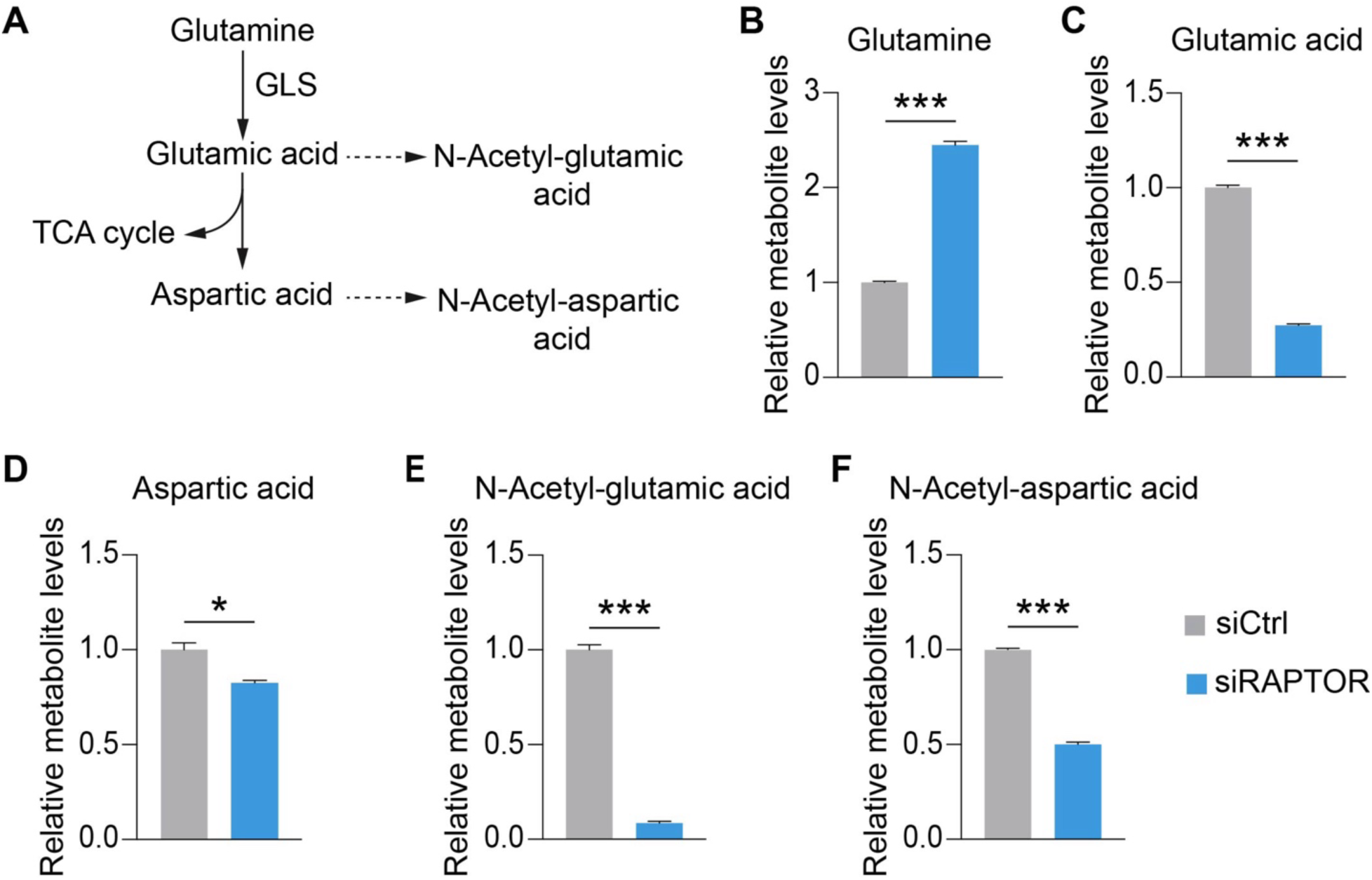
RAPTOR depletion impairs the generation of glutamine metabolism-derived metabolites in LECs. (A) Schematic illustrating the enzymatic pathway converting glutamine to aspartic acid. (B) Accumulation of glutamine in RAPTOR-depleted HDLECs compared with control HDLECs. (C, D) Reduced levels of glutamic acid and aspartic acid in HDLECs with RAPTOR knockdown. (E, F) Decreased levels of N-acetyl-glutamic acid and N-acetyl-aspartic acid in RAPTOR-deficient HDLECs. n = 3 samples for control siRNA-transfected HDLECs and n = 4 samples for RAPTOR siRNA-transfected HDLECs. Data represent mean ± SEM, *p < 0.05, ***p < 0.001, Welch’s t-test.

Glutamic acid can be converted to N-acetyl-glutamic acid, which functions as an essential activator of the urea cycle.^23^ Additionally, aspartic acid can be acetylated to form N-acetyl-aspartic acid in several cell types, particularly in neurons, where it has been implicated in diverse physiological processes ranging from gene expression regulation to brain metabolism.^24,25^ In our untargeted metabolomics studies, in addition to the core metabolites in glutaminolysis, we observed that N-acetyl-glutamic acid and N-acetyl-aspartic acid were also significantly reduced in RAPTOR-deficient HDLECs (**Fig. 2E and 2F**), indicating a broader impact of mTORC1 signaling on glutamine metabolism.

### Impaired glutaminolysis in RAPTOR-depleted LECs is accompanied by a compensatory increase in asparagine and upregulation of asparagine transporter expression

Asparagine is a non-essential amino acid (NEAA) that can be generated from aspartic acid. However, when glutaminolysis is impaired—such as under glutamine deprivation—glutamine-derived aspartate levels decline, limiting endogenous asparagine synthesis.^26,27^ Under these conditions, cells become dependent on extracellular asparagine, functionally rendering it an essential amino acid (EAA).^26,27^ Notably, extracellular asparagine has been shown to rescue cell proliferation and survival defects caused by glutamine deprivation.^26,27^ Given that our metabolomic analysis demonstrated that RAPTOR deficiency impaired glutaminolysis, we examined whether asparagine levels were altered. Interestingly, glutaminolysis impairment in RAPTOR-depleted HDLECs was accompanied by a marked increase in asparagine (**Fig. 3A**), potentially reflecting a compensatory response.

**Figure 3.**
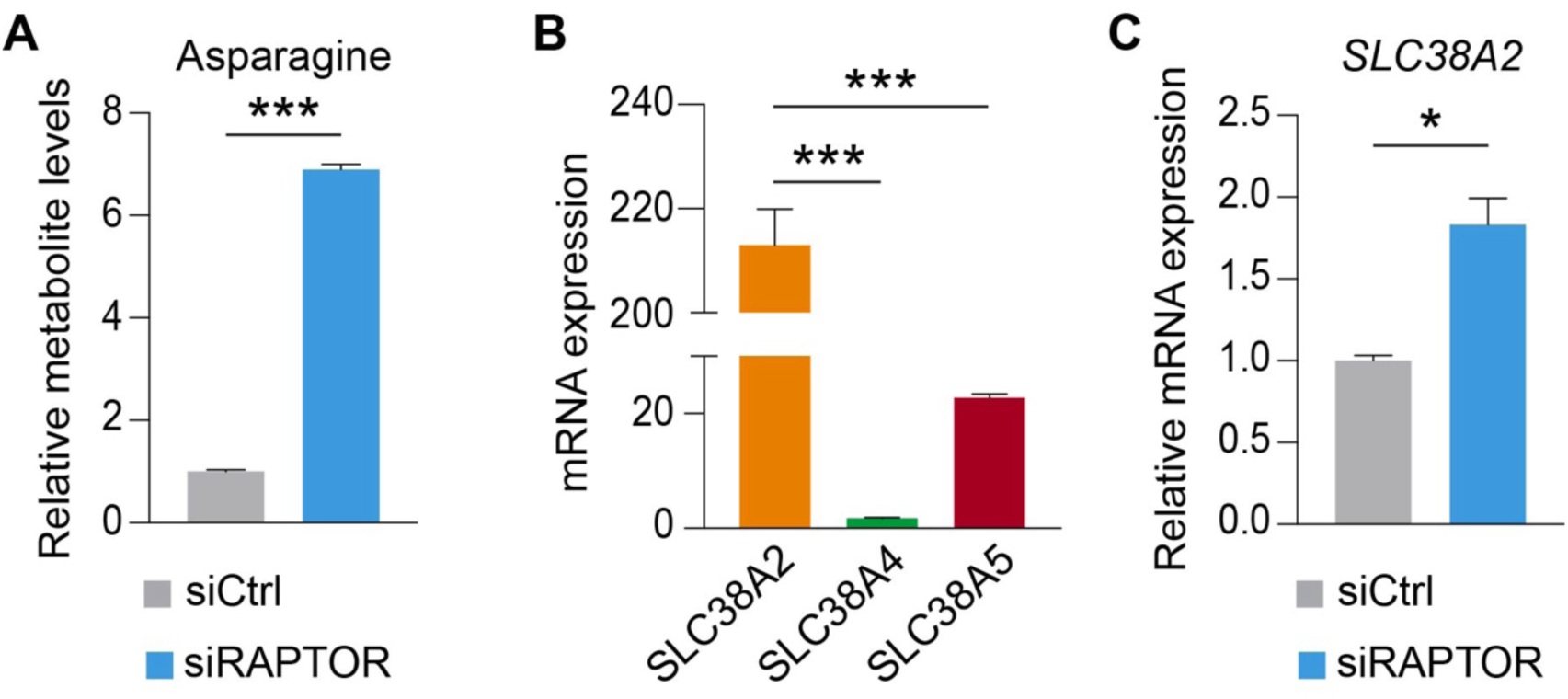
Compensatory increase in asparagine levels and upregulation of asparagine transporter expression in RAPTOR-deficient LECs. (A) Increased asparagine levels in RAPTOR-depleted HDLECs (n = 4 samples) compared with control HDLECs (n = 3 samples). (B) mRNA levels of three asparagine transporters in control HDLECs (n = 3 samples). (C) Upregulation of *SLC38A2* mRNA in RAPTOR-deficient HDLECs (n = 3 samples) compared with control HDLECs (n = 3 samples). Data represent mean ± SEM, *p < 0.05, ***p < 0.001, Welch’s t-test (A, C), or one-way ANOVA with Šídák’s multiple comparisons test (B).

We then asked how RAPTOR deficiency increased asparagine levels in HDLECs. To address this question, we focused on amino acid transporters involved in the transport of asparagine across the plasma membrane, which include SLC38A2, SLC38A4, and SLC38A5.^28^ Reanalysis of our previously published RNA sequencing (RNA-seq) dataset^18^ (Gene Expression Omnibus: GSE242082) indicated that SLC38A2 was the most abundant asparagine transporter in HDLECs (**Fig. 3B**). Notably, *SLC38A2* mRNA levels were increased by approximately 80% in RAPTOR-depleted HDLECs (**Fig. 3C**), suggesting that upregulation of *SLC38A2* may contribute to the elevated asparagine levels observed following RAPTOR knockdown.

### RAPTOR deficiency increases metabolites in arginine catabolism and arginine transporter expression in LECs

Arginine is synthesized primarily in the kidney and is classified as a semi-essential amino acid.^29^ Under physiological conditions, circulating arginine derived from renal production is sufficient to meet the cellular demands of most mammalian cell types.^30^ However, during infections, cancer, and other pathological conditions, circulating arginine may become insufficient, necessitating endogenous arginine synthesis to maintain intracellular arginine homeostasis.^30^

Arginine catabolism involves conversion of arginine to ornithine, which is subsequently metabolized into polyamines, including putrescine (**Fig. 4A**).^31^ Cellular polyamine levels need to be tightly regulated, as both deficiency and excessive accumulation can adversely affect cell proliferation or survival.^31^ Given the importance of arginine for cellular physiology, we quantified intracellular levels of arginine in our untargeted metabolomics dataset. We found that RAPTOR depletion resulted in significant elevation of arginine, ornithine, and putrescine in HDLECs (**Fig. 4B–4D**).

**Figure 4.**
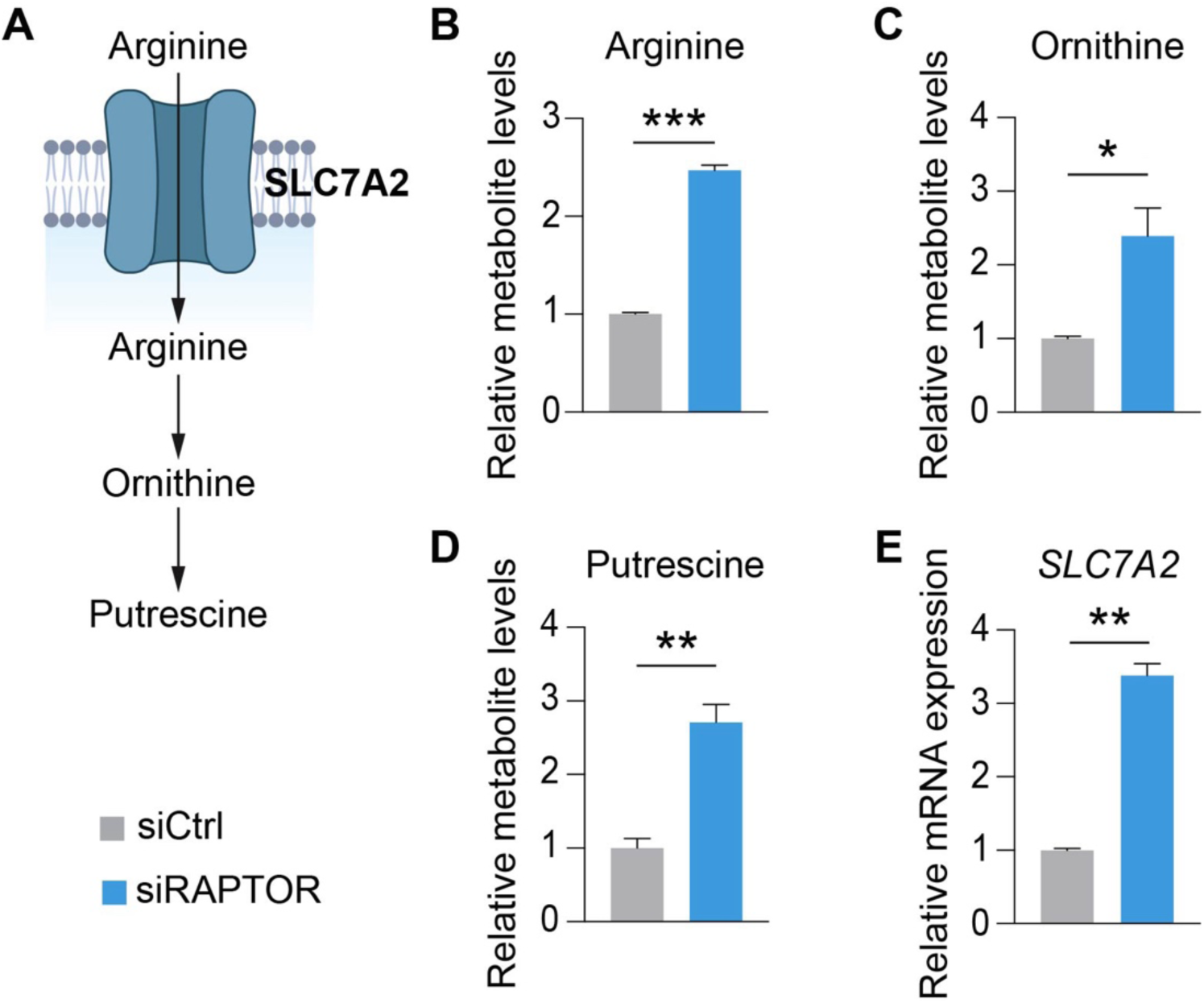
RAPTOR depletion increases metabolite levels in arginine catabolism and upregulates arginine transporter expression in LECs. (A) Schematic illustrating arginine import and catabolism. (B–D) Increased levels of arginine, ornithine, and putrescine in RAPTOR-deficient HDLECs (n = 4 samples) compared with control HDLECs (n = 3 samples). (E) Upregulation of *SLC7A2* mRNA levels in RAPTOR-depleted HDLECs (n = 3 samples) compared with control HDLECs (n = 3 samples). Data represent mean ± SEM, *p < 0.05, **p < 0.01, ***p < 0.001, Welch’s t-test.

To explore how RAPTOR deficiency may influence arginine metabolism, we examined whether arginine transporter expression in HDLECs was altered, given that most non-renal cells primarily acquire arginine from the extracellular environment under normal conditions. Key arginine transporters include cationic amino acid transporter (CAT)-1 (encoded by *SLC7A1*) and CAT-2 (encoded by *SLC7A2*). While CAT-1 is ubiquitously expressed, CAT-2 levels are highly regulated.^32^ We therefore analyzed our RNA-seq dataset and found that *SLC7A2* mRNA was increased by more than 2-fold in RAPTOR knockdown HDLECs (**Fig. 4E**), suggesting that arginine uptake is enhanced following mTORC1 inhibition.

### RAPTOR depletion impairs branched-chain amino acid (BCAA) catabolism in LECs

BCAAs, comprising valine, leucine, and isoleucine, are EAAs that play critical roles in metabolic homeostasis and have been implicated in the pathogenesis of cardiovascular diseases.^33,34^ For example, in a mouse model of heart failure with preserved ejection fraction (HFpEF), cardiac lymphatic vessels exhibit structural and functional abnormalities, which can be attributed to defective BCAA catabolism in cardiac LECs.^35^ Expression of enzymes involved in BCAA metabolism, including branched-chain amino acid transaminase 2 (BCAT2), which catalyzes the conversion of BCAAs to branched-chain α-keto acids, is significantly downregulated in cardiac LECs from mice with HFpEF.^35^ These alterations cause BCAA accumulation, resulting in impaired lymphangiogenesis.^35^

Given the important role played by BCAA catabolism in LEC function, we examined the impact of RAPTOR knockdown on intracellular BCAA levels. Metabolomic analysis revealed that valine and isoleucine concentrations were both increased by approximately 50% following RAPTOR depletion in HDLECs (**Fig. 5A and 5B**). Leucine was not detected in our untargeted metabolomics studies. To explore the potential underlying mechanism, we assessed *BCAT2* mRNA levels in our RNA-seq dataset from control and RAPTOR siRNA-transfected HDLECs and found that *BCAT2* expression was significantly reduced (**Fig. 5C**). These findings suggest that mTORC1 inhibition may impair BCAA catabolism in LECs.

**Figure 5.**
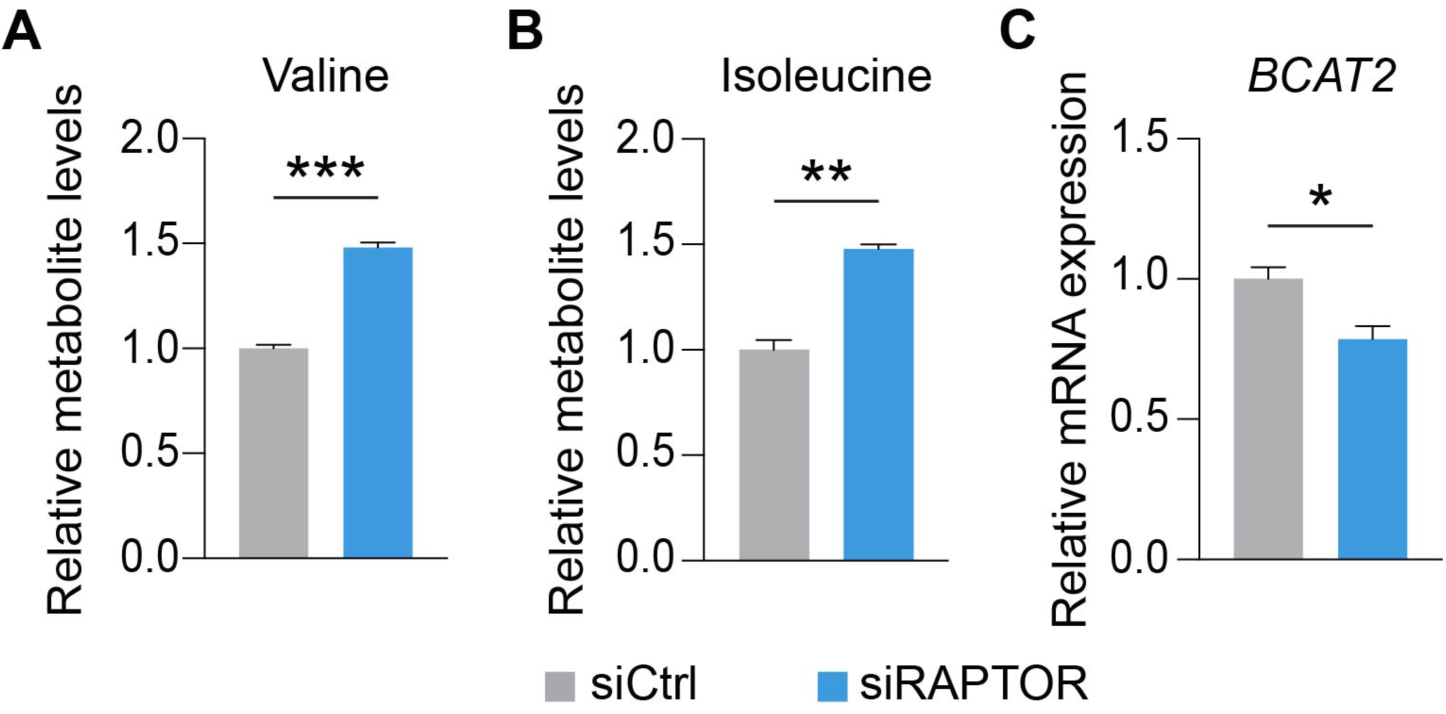
BCAA catabolism is impaired following RAPTOR depletion in LECs. (A, B) Increased levels of valine and isoleucine in RAPTOR-depleted HDLECs (n = 4 samples) compared with control HDLECs (n = 3 samples). (C) Downregulation of *BCAT2* mRNA levels in RAPTOR-deficient HDLECs (n = 3 samples) compared with control HDLECs (n = 3 samples). Data represent mean ± SEM, *p < 0.05, **p < 0.01, ***p < 0.001, Welch’s t-test.

### RAPTOR deficiency results in the accumulation of threonine, histidine, and lysine and impairs translational control in LECs

In addition to BCAAs, we quantified intracellular levels of other EAAs in our untargeted metabolomic analysis and found that threonine, histidine, and lysine were markedly accumulated in RAPTOR-depleted HDLECs (**Fig. 6A–6C**). The primary role of these EAAs is to support protein synthesis, as their catabolism is very limited in most mammalian cell types. For example, histidine catabolism begins with deamination catalyzed by histidine ammonia-lyase, which is encoded by the *HAL* gene.^36^ However, *HAL* was not expressed in HDLECs according to our RNA-seq dataset (data not shown). Similarly, lysine catabolism occurs predominantly in the liver.^37^

**Figure 6.**
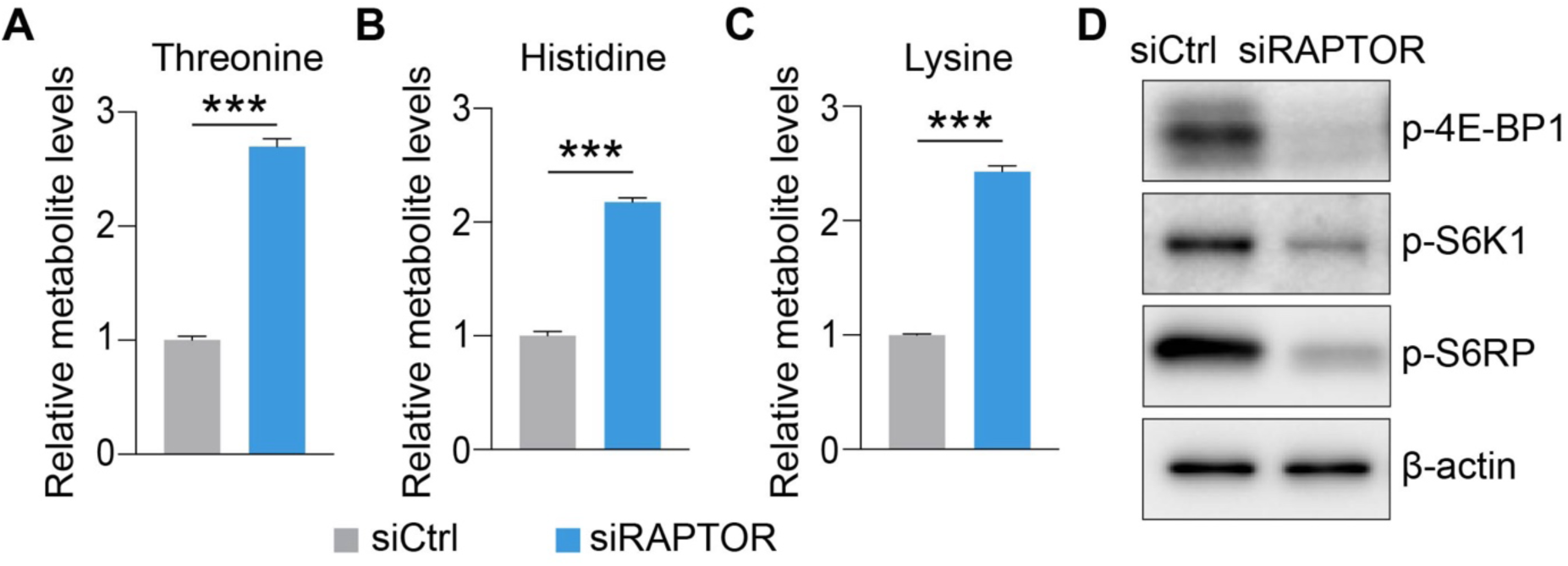
RAPTOR depletion leads to the accumulation of threonine, histidine, and lysine and impairs translational control in LECs. (A–C) Accumulation of threonine, histidine, and lysine in RAPTOR-deficient HDLECs (n = 4 samples) compared with control HDLECs (n = 3 samples). (D) Western blot analysis demonstrating that RAPTOR knockdown reduces phosphorylation of 4E-BP1, S6K1, and S6RP in HDLECs. Data represent mean ± SEM, ***p < 0.001, Welch’s t-test.

These observations suggest that the accumulation of threonine, histidine, and lysine in RAPTOR-deficient HDLECs is unlikely to result from impaired catabolism and instead may reflect reduced utilization of these EAAs for protein synthesis. We therefore focused on the impact of RAPTOR deficiency on translational control. To assess this, we examined the phosphorylation status of eukaryotic translation initiation factor 4E-binding protein 1 (4E-BP1) and the ribosomal protein S6 kinase 1 (S6K1)–S6 ribosomal protein (S6RP) axis, which are key downstream effectors of mTORC1 involved in the regulation of protein synthesis.^38^ Our results showed that phosphorylation of 4E-BP1, S6K1, and S6RP was markedly reduced following RAPTOR knockdown in HDLECs (**Fig. 6D**), suggesting suppressed translational activity.

## DISCUSSION

In this study, we performed untargeted metabolomic profiling to investigate the role of mTORC1 in LEC metabolism. Our data revealed an unexpected global alteration in amino acid levels, including NEAAs and EAAs, following RAPTOR knockdown in HDLECs. We recently discovered that mTORC1 promotes the formation of lymphatic capillaries and collecting vessels by regulating glycolysis and glutaminolysis.^18^ Specifically, RAPTOR deficiency downregulates HK2 and GLS expression and decreases TCA cycle metabolites.^18^ Consistent with these findings, our metabolomic analysis showed that isocitric acid levels are decreased in RAPTOR-depleted HDLECs. Moreover, glutamic acid and aspartic acid are reduced, whereas glutamine is accumulated, indicating that RAPTOR knockdown-induced GLS downregulation impairs the conversion of glutamine to glutamic acid. Notably, we also detected N-acetyl-glutamic acid and N-acetyl-aspartic acid, whose levels decreased following RAPTOR depletion in HDLECs.

Previous studies have reported that N-acetyl-glutamic acid is primarily enriched in the liver, whereas N-acetyl-aspartic acid is highly abundant in the mammalian brain.^39,40^ Therefore, the detectable abundance of these two N-acetylated metabolites in HDLECs was unexpected. In line with our observations, N-acetyl-aspartic acid has also been detected in adipocytes, where it influences lipid synthesis and gene expression through histone acetylation.^25,41^ These prior findings raise the possibility that N-acetyl-glutamic acid and N-acetyl-aspartic acid may play functional roles in HDLECs, which remains to be elucidated in future studies.

Previous studies in cancer cells have demonstrated that glutamine deprivation leads to asparagine depletion, resulting in proliferation defects and even apoptosis; however, these effects can be rescued by extracellular supplementation of asparagine.^26,27^ Interestingly, our untargeted metabolomic analysis revealed that RAPTOR deficiency-induced impairment of glutaminolysis is associated with elevated intracellular asparagine levels. This finding suggests that HDLECs may engage a compensatory mechanism to counteract the metabolic stress caused by mTORC1 inhibition. Consistent with this notion, our transcriptomic data further showed that the expression of *SLC38A2*, which encodes an asparagine transporter, is upregulated in RAPTOR-depleted HDLECs, thereby providing a potential mechanistic basis for the increased intracellular asparagine. Nevertheless, further studies are warranted to determine how mTORC1 regulates *SLC38A2* expression and whether simultaneous knockdown of RAPTOR and SLC38A2 results in more pronounced LEC proliferation defects compared with RAPTOR knockdown alone.

Impaired BCAA catabolism in LECs has been implicated in abnormal cardiac lymphatic vessel formation in HFpEF mice.^35^ In that context, reduced expression of BCAA-catabolizing enzymes in heart LECs leads to intracellular BCAA accumulation, which hinders vascular endothelial growth factor receptor 3 (VEGFR3) signaling and ultimately compromises lymphangiogenesis.^35^ Notably, our integrated metabolomic and transcriptomic analyses showed that RAPTOR deficiency results in increased levels of valine and isoleucine, accompanied by downregulation of *BCAT2*. These findings raise the possibility that RAPTOR deficiency-induced BCAA accumulation may impair VEGFR3 pathway activation in LECs, thereby contributing to defective lymphangiogenesis. Future studies will be required to determine whether altered BCAA metabolism directly mediates the lymphangiogenic defects observed in RAPTOR-depleted LECs and to elucidate the underlying molecular mechanisms.

Taken together, our work reveals a critical role of mTORC1 signaling in regulating amino acid uptake and catabolism in LECs. mTORC1 is well known to be activated by amino acids and thus plays a central role in nutrient sensing.^42,43^ Our study provides strong evidence for a reciprocal regulatory relationship between mTORC1 signaling and amino acid metabolism. A deeper understanding of this mutual regulation in LECs will provide important insights into the metabolic control of lymphangiogenesis.

## MATERIALS AND METHODS

### Cell Culture and Transfection

Human dermal lymphatic endothelial cells (HDLECs; brand name: HMVEC-dLyNeo-Der Lym Endo EGM-2MV; human lymphatic microvascular endothelial cells) were purchased from Lonza Bioscience. EBM-2 basal medium (Lonza Bioscience, CC-3156) with the EGM-2 MV SingleQuots Kit (Lonza Bioscience, CC-4147) was used for culturing HDLECs. Before seeding HDLECs, cell culture dishes were coated with 0.1% gelatin (Sigma-Aldrich) in a 37°C cell culture incubator to enhance cell attachment and then rinsed once with Dulbecco’s Phosphate-Buffered Saline (Thermo Fisher Scientific, 14190250). For gene knockdown experiments, HDLECs were transfected with small interfering RNAs (siRNAs) targeting RAPTOR (QIAGEN; Hs_KIAA1303_4, target sequence: TCGGACGTGGCCATGAAAGTA) or AllStars Negative Control siRNA (QIAGEN), which was used as a non-targeting control. siRNA transfections were performed using Lipofectamine RNAiMAX reagent (Thermo Fisher Scientific).

### Untargeted LC-MS Metabolomics

Metabolites in HDLECs were extracted using an 8:2 (v/v) methanol:H₂O solution pre-chilled to −75 °C. The untargeted LC-MS metabolomics method adopted in this work has been used in our published studies.^44–50^ Briefly, all LC-MS experiments were conducted using a Thermo Vanquish UPLC-Exploris 240 Orbitrap MS instrument (Waltham, MA). Each sample was injected twice, 10 μL for analysis using negative ionization mode and 4 μL for analysis using positive ionization mode. Both chromatographic separations were performed in hydrophilic interaction chromatography (HILIC) mode on a Waters XBridge BEH Amide column (150 x 2.1 mm, 2.5 µm particle size, Waters Corporation, Milford, MA). The flow rate was 0.3 mL/min, the auto-sampler temperature was kept at 4 °C, and the column compartment was set at 40 °C. The mobile phase was composed of Solvents A (10 mM ammonium acetate, 10 mM ammonium hydroxide in 95% H_2_O/5% ACN) and B (10 mM ammonium acetate, 10 mM ammonium hydroxide in 95% ACN/5% H_2_O). After the initial 1 min isocratic elution of 90% B, the percentage of Solvent B decreased to 40% at t = 11 min. The composition of Solvent B was maintained at 40% for 4 min (t = 15 min), and then the percentage of B gradually went back to 90% to prepare for the next injection. Using a mass spectrometer equipped with an electrospray ionization (ESI) source, we collected untargeted data from 70 to 1050 m/z.

To identify peaks from the MS spectra, we made extensive use of the in-house chemical standards (approximately 600 aqueous metabolites). Additionally, we searched the resulting MS spectra against the HMDB library, Lipidmap database, METLIN database, and commercial databases, including mzCloud, Metabolika, and ChemSpider. The absolute intensity threshold for the MS data extraction was 1,000, and the mass accuracy limit was set to 5 ppm. Identifications and annotations used available data for retention time (RT), exact mass (MS), MS/MS fragmentation pattern, and isotopic pattern. We used the Thermo Compound Discoverer 3.3 software for aqueous metabolomics data processing. The untargeted data were processed by the software for peak picking, alignment, and normalization. To improve rigor, only the signals/peaks with CV < 20% across quality control pools and the signals showing up in >80% of all the samples were included for further analysis.

### Western blotting

Total cellular protein was extracted using a radioimmunoprecipitation assay (RIPA) lysis buffer, and protein concentration was quantified by the bicinchoninic acid (BCA) assay. Equal amounts of protein were loaded and resolved on SDS-polyacrylamide gels and subsequently transferred to polyvinylidene difluoride (PVDF) membranes. Membranes were incubated in 5% nonfat milk for 45 minutes at room temperature to reduce nonspecific binding, followed by overnight incubation at 4°C with primary antibodies. After washing, membranes were incubated with horseradish peroxidase-conjugated secondary antibodies. Chemiluminescent signal detection was performed using SuperSignal™ West Pico PLUS chemiluminescent substrate or SuperSignal™ West Femto Maximum Sensitivity Substrate (Thermo Fisher Scientific), and images were captured using a ChemiDoc MP system (Bio-Rad). Primary antibodies included RAPTOR (Cell Signaling Technology, 2280), phospho-4E-BP1 (Cell Signaling Technology, 2855), phospho-S6K1 (Cell Signaling Technology, 9205), phospho-S6 ribosomal protein (Cell Signaling Technology, 4858), and β-actin (Sigma-Aldrich, A5316). Band density was quantified using ImageJ.

### Statistical and data analysis

Statistical analyses were performed in R (version 2025.05.1+513). Given the small sample size (n = 3–4 per group), parametric testing was selected because the t-test is generally robust to moderate deviations from normality and provides greater statistical power than nonparametric alternatives in small datasets.^51^ To account for multiple comparisons across metabolites, p-values were adjusted using the false discovery rate (FDR) method (Benjamini–Hochberg procedure). Metabolites with an adjusted p-value (padj) < 0.05 were considered statistically significant. To evaluate overall metabolic differences and assess sample clustering, PCA was performed using the PRcomp function in R on scaled data. PCA results were visualized using the autoplot function. Principal component 1 (PC1) explained 50.89% of the total variance, while PC2 accounted for approximately 18%. Samples segregated according to experimental groups, indicating distinct metabolic profiles. For the other statistical comparisons, Welch’s t-test or one-way ANOVA with Šídák’s multiple comparisons test were used to determine statistical significance.

## ACKNOWLEDGEMENTS

We thank the OMRF Center for Biomedical Data Sciences for assistance with the statistical analysis of the untargeted metabolomics data. P.Y. is supported by grants from the National Institutes of Health (HL162985 and GM139763), the American Heart Association (23IPA1054589 and 25TPA1481189), OMRF internal research funding, the Oklahoma Center for Adult Stem Cell Research, and the Presbyterian Health Foundation.

